# Lyb-2-4, a unique genomic region of hypervirulent carbapenem-resistant *Acinetobacter baumannii*

**DOI:** 10.1101/768655

**Authors:** Chao Zheng, Jinyong Zhang, Min Zhang, Zhou Liu, Yuxin Zhong, Shiyi Liu, Ruiqin Cui, Qiuyang Deng, Yanji Xu, Yun Shi, Hao Zeng, Xiyao Yang, Chuchu Lin, Yutian Luo, Huaisheng Chen, Weiyuan Wu, Jinsong Wu, Tianle Zhang, Yuemei Lu, Xueyan Liu, Quanming Zou, Wei Huang

## Abstract

*Acinetobacter baumannii* is an important human pathogen due to its multi-drug resistance, but is usually with low-grade virulence. Although a mouse model revealed different virulence grades of clinical carbapenem-resistant *A. baumannii* (CRAB) strains, the genetic basis remains unknown. We collected 61 CRAB isolates from intensive care unit of Shenzhen People’s Hospital (Shenzhen, China), and analyzed them used whole genome sequencing (WGS), multilocus sequence typing (MLST) and core genome MLST (cgMLST), transmission chain reconstruction and Comparative genomic tools. A mouse pneumonia model was used to confirm the hypervirulent phenotype. Eleven complex types (CT) were identified based on core genome multilocus sequence typing scheme. CT512 showed higher transmissibility and bloodstream infection rates than other CTs. A genomic region Lyb-2-4 was shared by CT512 and CT2092 but not CT2085. The mortality rates of patient infected with CRAB harboring Lyb-2-4 was significantly higher than those infected with CRAB isolates without Lyb-2-4 (77.8% vs 24.5%, *p* < 0.01). In the mouse model, the survival rates of strains containing the Lyb-2-4 region (LAC-4, 5122 and 2092) were significantly lower than for strains without Lyb-2-4 (7152, 71517, 20859 and ATCC17978). One open reading frame (ORF) was a marker for the presence of Lyb-2-4, and PCR of a segment of this ORF, designated as *hvcT*, served as a tag for hypervirulent CRAB. Our study should be very useful in advising the clinician to implement medical intervention earlier, and also making the worldwide surveillance of these hypervirulent CRAB strains easier.

**IMPORTANCE:** Hypervirulent CRAB strains are expected to pose a threat to human health because infection of these strains is associated with high mortality and multidrug resistance. The rapid hypervirulent CRAB identification assay will facilitate prompt medical intervention. Our findings should provoke surveillance for hypervirulent CRAB strains harboring Lyb-2-4 in other countries. Further research should focus on the mechanism of hypervirulence, the acquisition of this genomic region and the development of control measures to prevent further dissemination.

---

*Acinetobacter baumannii* is a gram-negative coccobacillus that can cause serious infections among critically ill patients, particularly in the intensive care unit (ICU) setting (1). Worldwide, *A. baumannii* accounted for up to 20% of infections in ICUs (2).

*A. baumannii* is recognized as one of the most problematic bacterial pathogens due to its propensity to acquire multidrug, extensive drug and even pan-drug resistance phenotypes (3). The highly efficacious and low toxicity carbapenems are regarded as important antibiotics for treating

*A. baumannii* infections (2), and the emergence and rapid spreading of carbapenem-resistant *A. baumannii* (CRAB) isolates are a global concern because they pose a severe threat to public health (4). WHO has included CRAB in the critical group in the list of bacteria that pose the greatest threat to human health, prioritizing research and development efforts for new antimicrobial treatments (5).

In addition to the rapid development of resistance to multiple antibiotics, virulence is also a noteworthy feature of *A. baumannii*. In general, *A. baumannii* has been regarded as a low-grade pathogen. Most laboratory strains and clinical isolates do not cause severe infections in immunocompetent mice, inducing only a self-limiting pneumonia with very limited local bacterial replication and systemic dissemination, even when a large inoculum is used (6, 7). However, studies have shown that some clinical CRAB strains are lethal to immunocompetent mice and thus revealed the range of virulence in different strains of this pathogen (8, 9). LAC-4 is a hypervirulent strain that killed 100% of mice within 48 hours after being inoculated with 10^8^ colony-forming units (CFU) (9). Even though the complete genome has been sequenced, the genetic basis of the hypervirulent phenotype is still not clearly understood (10).

Multilocus sequence typing (MLST) has been the most commonly used technique for defining *A. baumannii* populations. Patients with Pasteur scheme sequence type 2 (ST2) bloodstream infections were less likely to be treated with appropriate empirical antibiotics and had higher mortality than those with non-ST2 bloodstream infections (11). Now, however, with fast and affordable whole genome sequencing (WGS), it is possible to compare whole genomes for strain typing rather than just the few loci used in traditional MLST. Genotyping with core genome MLST (cgMLST) uses thousands of alleles across the genome, therefore resulting in a higher level of strain discrimination. The cgMLST scheme for *A. baumannii* was developed and evaluated in 2017 (12).

In this study, we perform a retrospective molecular epidemiological study to clarify the population composition of CRAB in the ICU of the Shenzhen People’s Hospital. By integrating cgMLST with clinical data we identified a sub-linkage of ST2 with high transmissibility and mortality that is emerging as an endemic strain in the hospital. Through the use of comparative genomics tools we also identified a genomic region, Lyb-2-4, that is associated with high mortality CRAB infections. We then developed a rapid tool to detect the presence of Lyb-2-4, which has the potential to be a novel marker of hypervirulent CRAB. Finally, we confirmed the hypervirulent phenotype in a mouse infection model.

## RESULTS

Non-repetitive CRAB (n=61) isolates collected from patients with confirmed *A. baumannii* infections were frozen and stored by the clinical microbiology laboratory at the Shenzhen People’s Hospital. Among all confirmed CRAB strains, 100% carried the OXA-23 carbapenemase and 1 strain carried both OXA-23 and KPC-2 (FIG. S1).

Classical Pasteur scheme MLST analysis identified all but one of the 61 CRAB isolates as belonging to the ST2 lineage, with the remaining isolate belonging to ST221. In contrast, cgMLST identified 11 CTs. The most common were CT715 (18 isolates: 7151 to 71518), CT2082 (14 isolates: 20821 to 208214), and CT2085 (10 isolates: 20851 to 208510), followed by CT512 (8 isolates: 5121 to 5128), CT2088 (3 isolates: 20881 to 20883), and CT2094 (3 isolates: 20941 to 20943). Five other CT types were identified in single isolate (Fig 1). Virulence gene analysis showed that most isolates shared the same profile except: *abaR* was missing in isolates 5121, 5122, 5123, 5127, 2096, 2092 and 2093; *bap* was missing in isolate 2096; and *hemO* was missing in isolates 20881 and 20883 (FIG. S2).

**FIG 1.**
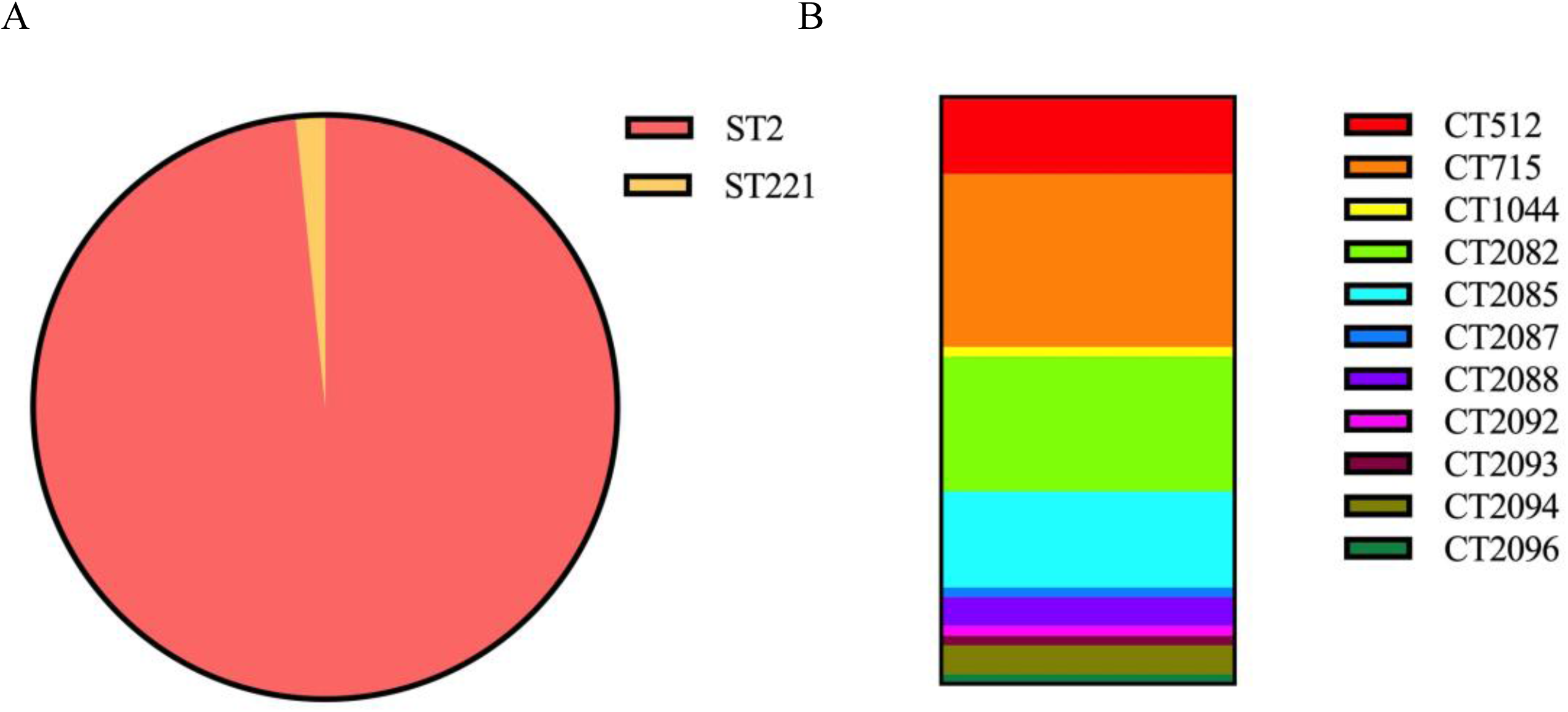
Distribution of 61 carbapenem-resistant *Acinetobacter baumannii* genotype in this study. (A) Genotype distribution of the isolates based on MLST scheme and (B) cgMLST scheme. ST=sequence type. CT=complex type.

We constructed minimum spanning trees (MST) with aim to identify the relationships among the different CTs. Five clusters were separated by a mean pairwise allelic distance of 9 alleles (FIG. 2B). Throughout the isolation timeline in 2017, the CTs appeared in sequence (FIG. 2A). CT2085 was prevalent during the first half of year while CT2082 and CT2088 were prevalent during the second half of year, and CT715 appeared in the transition time between the first and second half of the year. The position of CT2082 on the MST was consistent with its possible role as a progenitor of CT715, while the position of CT715 was consistent with its possible role as a progenitor of CT2082 and CT2088. The phylogenetic relationships were consistent with the isolation timeline, making it possible to trace the direction of evolution.

**FIG 2.**
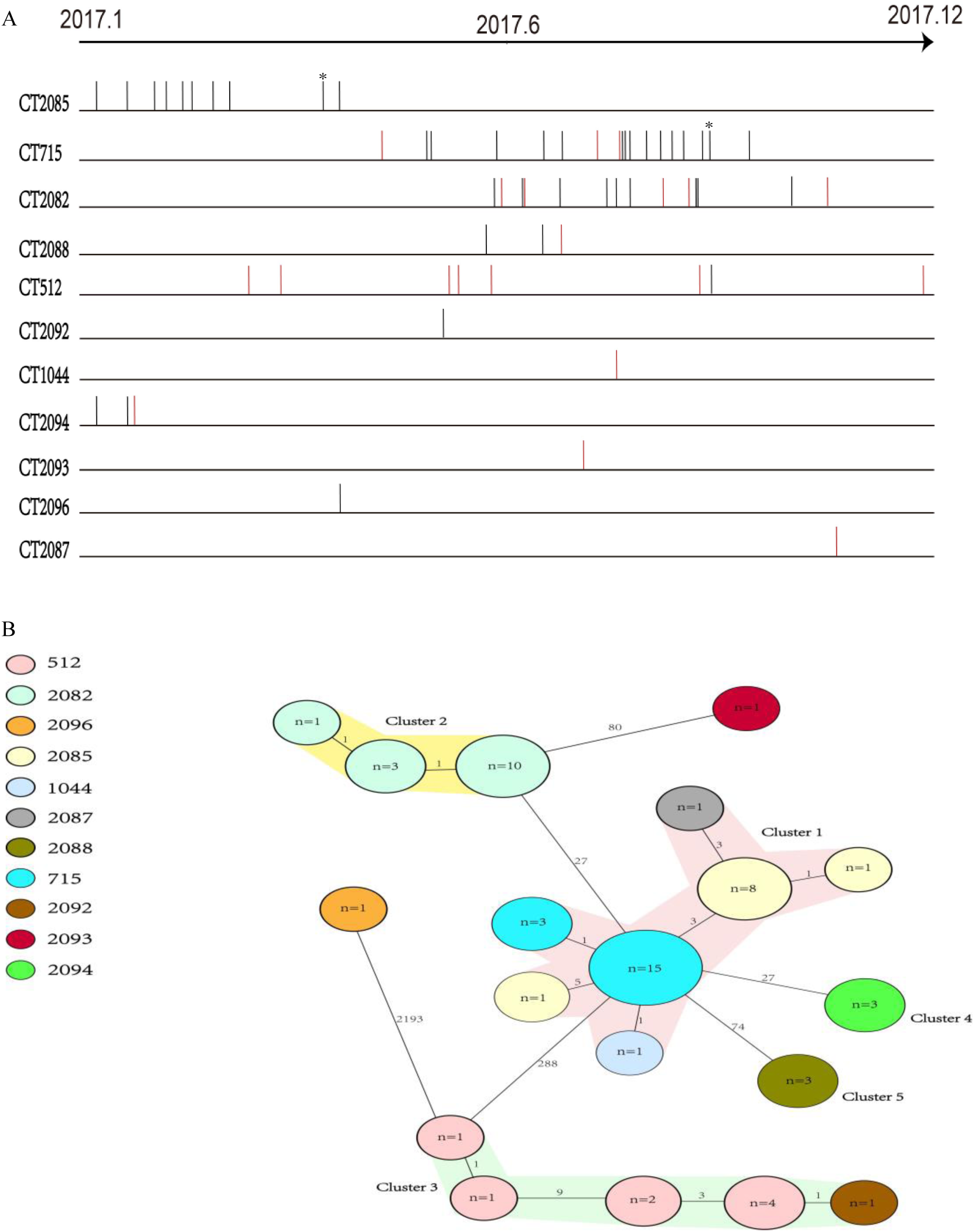
Timelines and minimum spanning trees of carbapenem-resistant *Acinetobacter baumannii* in this study. (A) Sampling dates of each isolate. Vertical lines represent the isolates in each CT group and red vertical lines represent death cases. * Some genes in Lyb-2-4 scattered presence in the genome of isolates. (B) Minimum spanning tree analysis of carbapenem-resistant *A. baumannii*. Each circle represents isolates belonging to a corresponding CT type based on sequence analysis of 2390 cgMLST target genes. The codes in the circles refer to case number. The numbers on the branches indicate the number of alleles distance. Cluster distance threshold is 9.

We integrated the epidemiological and WGS data to construct transmission trees. We found that the probability of transmission events within CT512 were extremely high (FIG. 3). Analysis of the clinical characteristics of the 61 patients with CRAB infections (table 1) showed that CT512 was associated with significantly higher mortality and bloodstream infection rates than the other CTs (Table 1). There were no statistically significant differences between the CTs in terms of demographics, types of admission, infection types, and main comorbidities.

**FIG 3.**
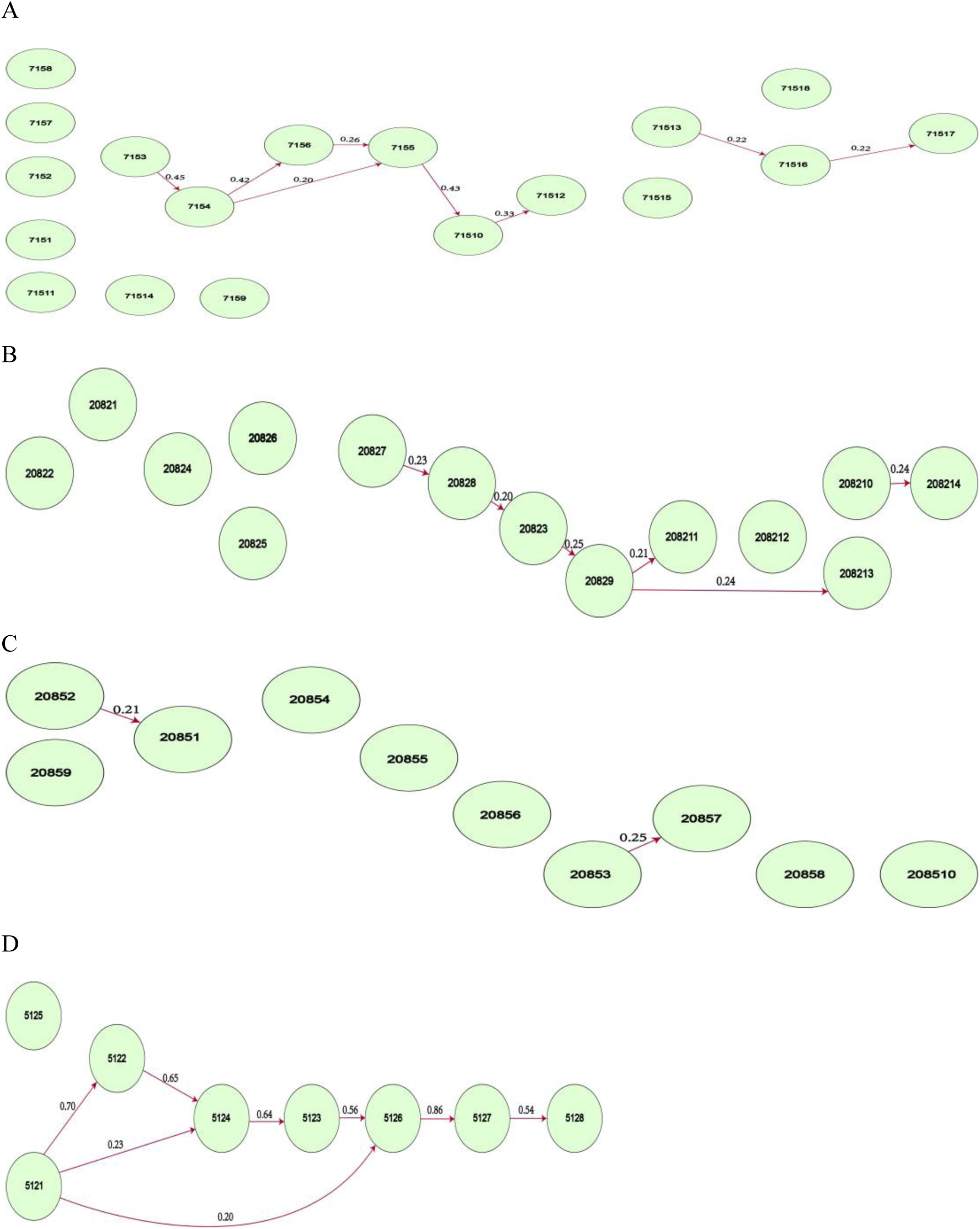
The transmission network of CTs with more than 3 cases. (A) CT715; (B) CT2082; (C) CT2085; (D) CT512; Number on each arrow represents the inferred probability of the corresponding transmission event. CT=complex type.

**Table 1:** Demographic and clinical characteristics of 61 patients with carbapenem-resistant *Acinetobacter baumannii* infections in this study.

|  | CT512<br>(n=8) | CT715<br>(n=18) | CT2082<br>(n=14) | CT2085<br>(n=10) | Other CTs<br>(n=11) | p value* |
| --- | --- | --- | --- | --- | --- | --- |
| Age in years, mean ± SD | 53.3 ± 23.6 | 59.9 ± 18.1 | 53.1 ± 20.9 | 64.6 ± 11.6 | 63.7 ± 18.4 | 0.444 |
| Gender (Male) | 5(62.5%) | 15(83.3%) | 10(71.4%) | 9(90.0%) | 7(63.6%) | 0.444 |
| Bacteremia | 3(37.5%) | 0(0.0%) | 1(7.2%) | 0(0.0%) | 2(18.2%) | **0.024** |
| Pulmonary infection | 4(50.0%) | 14(77.8%) | 11(78.6%) | 9(90.0%) | 6(54.5%) | 0.185 |
| Urinary infection | 0(0.0%) | 1(5.6%) | 0(0.0%) | 0(0.0%) | 0(0.0%) | 0.647 |
| Abdominal infection | 0(0.0%) | 3(16.7%) | 2(14.3%) | 1(10.0%) | 4(36.4%) | 0.257 |
| Cerebral vascular disease | 5(62.5%) | 6(33.3%) | 7(50.0%) | 6(60.0%) | 4(36.4%) | 0.421 |
| Hypertension | 4(50.0%) | 9(50.0%) | 7(50.0%) | 6(60.0%) | 7(63.6%) | 0.957 |
| Diabetes | 1(12.5%) | 4(22.2%) | 1(7.1%) | 3(30.0%) | 4(36.4%) | 0.432 |
| Mechanical ventilation<sup>SEP</sup> | 8(100%) | 16(88.9%) | 14(100.0%) | 8(80.0%) | 9(81.8%) | 0.190 |
| Central venous catheter | 6(75.0%) | 15(83.3%) | 10(71.4%) | 7(70.0%) | 9(81.8%) | 0.771 |
| Foley catheter | 8(100.0%) | 17(94.4%) | 13(92.9%) | 10(100%) | 11(100.0%) | 0.422 |
| Nasogastric tube | 6(75.0%) | 12(66.7%) | 13(92.9%) | 7(70.0%) | 11(100.0%) | 0.097 |
| Continuous renal replacement therapy | 3(37.5%) | 5(27.8%) | 4(28.6%) | 3(30.0%) | 5(45.5%) | 0.852 |
| Mortality | 7(87.5%) | 3(16.7%) | 5(35.7%) | 0(0.0%) | 4(36.4%) | **0.001** |

CT=complex type. \* p values are for comparisons between CTs. Dichotomous variables are analyzed using chi-square test and continuous variables are analyzed using one-way ANOVA.

High transmissibility and mortality rate highlighted the distinctiveness of CT512. To try to identify the genetic basis of the more virulent phenotype of the CT512 strains, we compared the genomes of CT512 strains with the genomes of CT2085 strains, because no deaths were associated with the strains belonging to CT2085. A gene cluster was identified that is shared by all CT512 genomes and absent from all CT2085 genomes. There appeared to be a high mortality associated with strains containing this special genomic region, which we designated Lyb-2-4. This is a region of 18813 bp containing 12 genes encoding proteins of known function and 7 genes whose putative encoded proteins have unknown functions (FIG. 4A; Table 2). Lyb-2-4 contains a series of genes (*leg1, leg2, leg4, leg5* and *legF*) encoding enzymes necessary for the biosynthesis of legionaminic acid, the precursor for an uncommon sugar (α-8-epi-legionaminic acid) found in the repeating unit of the surface polysaccharide of *A. baumannii* (13, 14). The *pglD* encodes an acetyltransferase for the biosynthesis of UDP-N,N’-diacetylbacillosamine (UDP-diNAcBac), a unique carbohydrate produced by a number of bacterial species, and which has been implicated in pathogenesis (15). The *fnlA, fnlB* and *fnlC* gene cluster in Lyb-2-4 encodes proteins that catalyze the biosynthesis of α-L-frucosamine, the precursor sugar for the biosynthesis of the surface polysaccharide of hypervirulent strain LAC-4 (13). The *wbuB* gene encodes an N-acetyl-L-fucosamine (L-FucNAc) transferase. The *gnu* encodes an epimerase probably involved in the biosynthesis of glucosamine. The product of *wbpL* catalyzes the formation of a phosphodiester bond between a membrane-associated undecaprenyl-phosphate (Und-P) molecule and N-acetylhexosamine 1-phosphate, which is usually donated by a soluble UDP-N-acetylhexosamine precursor. In addition, Lyb-2-4 contains a series of genes with unknown functions (*orf1*-*orf7*). The encoded protein of *orf1* shares 85% identity with the protein Leg6 that is involved in the biosynthesis of legionaminic acid. The genes of *orf2* and *orf3* encode putative proteins belonging to different protein families of oxidoreductases. The products of *orf6* and *orf7* are probably involved in the pathway of polysaccharide biosynthesis, and the functions of *orf4* and *orf5* are completely unknown. We scanned the genome of the other CT types and found that Lyb-2-4 was also found in the isolate 2092. In addition, some genes present in Lyb-2-4 were found scattered around the genomes of isolates 71517 and 20859 (Fig. 2A).

**FIG 4.**
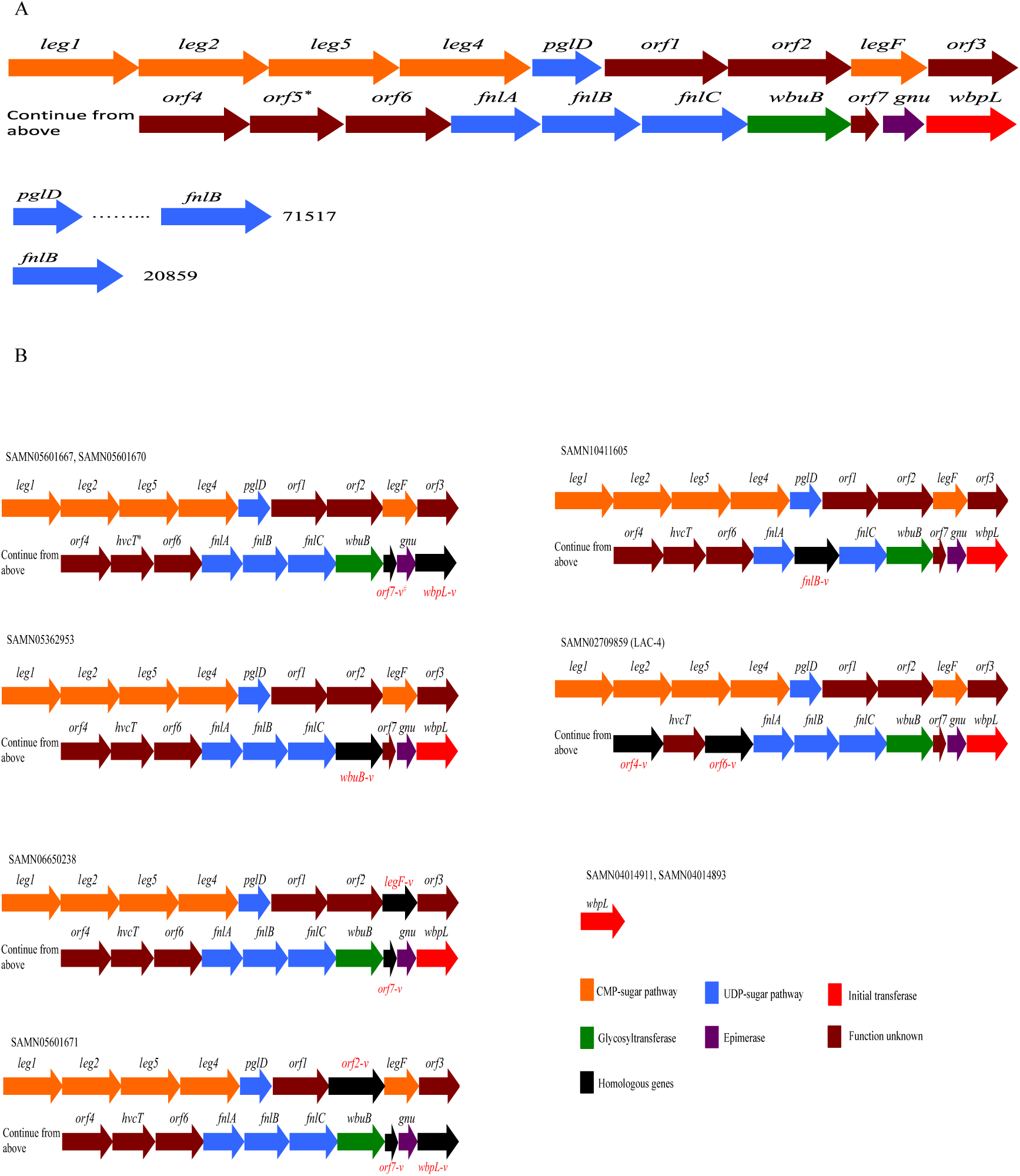
Gene organization and functional assignment in Lyb-2-4. (A) Organization of Lyb-2-4 and some genes scattered presence in the strains of this study. (B) Organization of Lyb-2-4 variants by scanning 140 complete genomes of *Acinetobacter baumannii* from NCBI. * We kept *orf5* as hypervirulent carbapenem-resistant *Acinetobacter baumannii* (CRAB) marker candidate and designated hypervirulent CRAB tag (*hvcT*). ^#^The presence of homologous genes in Lyb-2-4 variants.

**Table 2:**
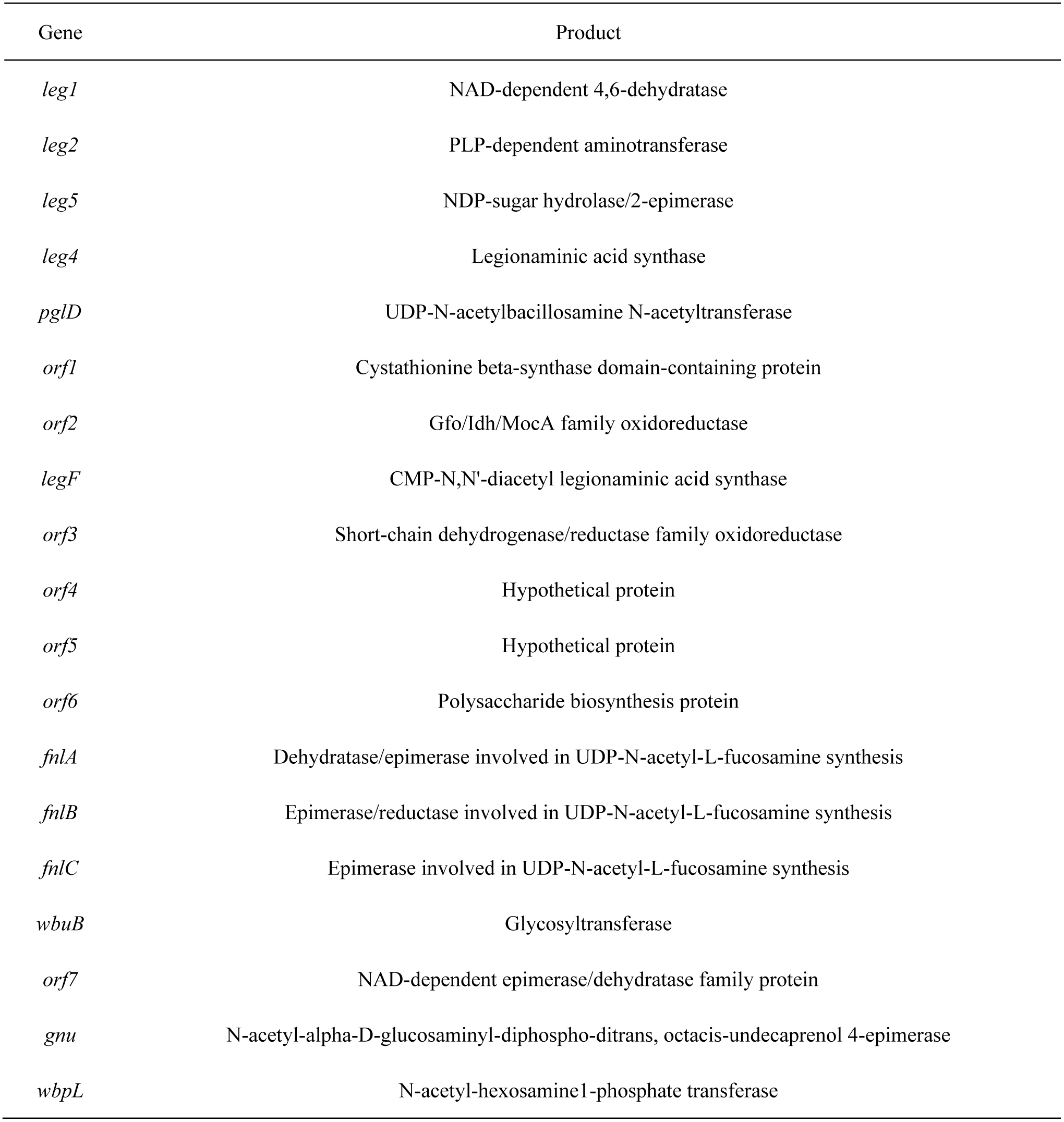
The products of genes in Lyb-2-4.

The isolates 5122, 7152, 71517, 20859, 2092, LAC-4 and ATCC17978 were selected to test their serum resistance and virulence in the mouse model (Fig. 4A). With an inoculum of 2.5×10^8^ CFU, the survival with isolates 5122, 2092 and LAC-4 was 0% at 48 hours, and at 7 days was 60% with ATCC17978, 50% with 7152, 71517 and 20859. With an inoculum of 5×10^7^ CFU, survival was 0% with LAC-4 at 48 hours; 10% with 5122 and 2092, 80% with ATCC17978 and 7152, 60% with 71517, and 90% with 20859 at 7 days. With an inoculum of 1×10^7^ CFU, 7 days survival was 10% with LAC-4, 40% with 5122, 30% with 2092, 90% with 71517, 100% with ATCC17978 and 7152. The survival rates of Lyb-2-4 harboring isolates (LAC-4, 5122 and 2092) were significantly lower than isolates not carrying Lyb-2-4 (7152, 71517, 20859 and ATCC17978) (FIG. 5; *p* < 0.0001 by log-rank test). No statistically significant differences of survival in 20% serum (FIG. S3).

**FIG 5.**
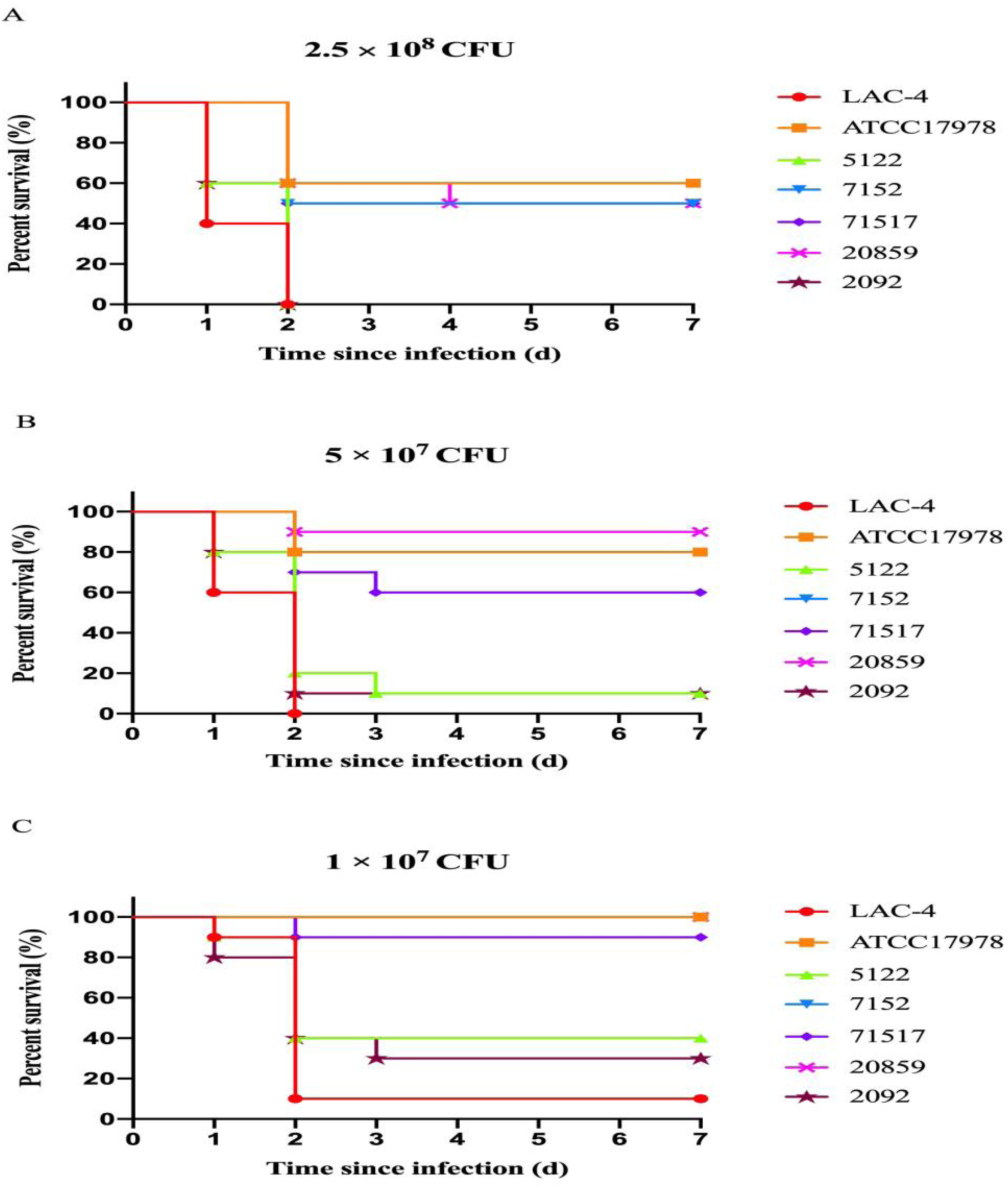
Virulence potential of *Acinetobacter baumannii* strains in a mouse infection model. The effect of 2.5×10 ^8^, 5×10 ^7^, 1×10 ^7^ colony-forming units of each *A. baumannii* isolate on survival was assessed in mouse. ATCC17978, 7152, 71517 and 20859 are *A. baumannii* isolates that did not carry Lyb-2-4. LAC-4 is a ST10 hypervirulent *A. baumannii* strain reported in a previous study. Isolates 5122 and 2092 are two ST2 *A. baumannii* strains harboring Lyb-2-4 in this study.

We browsed 140 complete genomes of *A. baumannii* from NCBI (Table S2) and found 2 strains (SAMN03817045 and SAMN03890258) that harbored Lyb-2-4; 7 strains, including LAC-4, that harbored a region homologous to Lyb-2-4 (FIG 4B); and 2 strains that only harbored *wbpL*. Based on our analysis, there are 13 genes linked with the presence of Lyb-2-4, but not *pglD, legF, orf4, wbuB, fnlB* or *wbpL*. We retrospectively used PCR to detect these 13 genes in 61 strains, and then filtered out 12 through specificity validation. Finally, we chose *orf5* as a hypervirulent CRAB marker candidate and identified a hypervirulent CRAB tag that we designated as *hvcT*. BLASTN to search the NCBI sequence databases verified that only *A. baumannii* strains harboring Lyb-2-4 possess *hvcT* sequence. We then used forward primer (5’-TTGAGAAGCTAAATTATGGCTCG-3’) and reverse primer (5’-GATAGCACAGAAATCCATAAAGGAA-3’) to screen for the presence of *hvcT*, as a marker for the presence of the hypervirulent locus Lyb-2-4. The supplementary data shows the sequence of Lyb-2-4, homologous genes in Lyb-2-4 variants and amplification conditions of *hvcT*.

For a retrospective investigation to confirm the correlation between infection with *hvcT*^*+*^ CRAB sisolates and mortality, we collected CRAB isolates (n=227) from 3 provinces of China, along with information on the clinical outcome. The mortality rates in patients infected with *hvcT*^*+*^ CRAB strains were significantly higher than for patients infected with *hvcT*^*-*^ CRAB strains (Table 3; 57.1% vs 20.8%, *p* < 0.0001 by chi-square test).

**Table 3:** Correlation between *hvcT*^*+*^ to high mortality.

| | $hvcT^+$ | $hvcT^-$ |
| --- | --- | --- |
| Death | 20 | 40 |
| Non-death | 15 | 152 |

$hvcT$ =hypervirulent carbapenem-resistant *Acinetobacter baumannii* tag. $p < 0.0001$ , p value is for mortality comparison between $hvcT^+$ and $hvcT^-$ strains using chi-square test.

## DISCUSSION

Our study investigated the population composition and evolutionary history of CRAB strains isolated in a Chinese hospital over one year period, using the MLST, cgMLST scheme and phylogenetic relationships. We demonstrated an association between infection with CRAB strains harboring genomic region Lyb-2-4 and high mortality rates in the ICU, and thus highlight the serious threat posed by CRAB strains harboring Lyb-2-4. We also developed a tool for the rapid identification of strains harboring Lyb-2-4 that could facilitate the identification of hypervirulent strains without the need for WGS, thereby providing an early warning so clinicians can implement adequate treatment promptly.

Virulence factors, such as secretion systems, surface glycoconjugates and micronutrient acquisition systems, facilitate *A. baumannii* pathogenesis. Capsular polysaccharide, glycosylated proteins, lipopolysaccharide and peptidoglycan are the common bacteria glycoconjugates. For *A. baumannii*, lipooligosaccharide (LOS) is the primary component of the outer leaflet of the outer membrane and as such is generally considered an essential structural component that is required for bacterial viability. Most *A. baumannii* strains also carry a thick capsular polysaccharide responsible for protecting the bacteria from external threats, including host defences (16). The products of the genes in Lyb-2-4 appear to be mostly involved with the synthesis of surface glycoconjugates: Lyb-2-4, *pglD, legF, fnlA, fnlB, fnlC, wbuB, gnu* and *wbpL* encode proteins known to be involved in polysaccharide biosynthesis, and the peptides encoded by *orf2, orf3, orf6* and *orf7* are putatively also involved in the capsular polysaccharide biosynthesis pathway (table 2). In addition, Similar to GI12 of LAC-4, Lyb-2-4 harbors a series of genes (*leg1, leg2, leg4, leg5, legF and leg6* homologue *orf1*) encoding enzymes necessary for the biosynthesis of legionaminic acid, and bacterial pathogens may utilize this unusual sugar to mimic the host cell surface, thus escaping from host immune surveillance and thereby facilitating colonization and invasion (13, 14).

Horizontally acquired genes are typically found grouped together in blocks of sequence called pathogenicity islands, fitness islands, or more generally, genomic islands (GI) (17). The investigation of genomic islands in *A. baumannii* found 12 in the genome of the hypervirulent strain LAC-4, including the 33.4-kb GI12 that contains a large number of genes involved in capsule biosynthesis (10). The structure of the polysaccharide isolated from LAC-4 showed that the polysaccharide was built of tri-saccharide repeating units containing α-L-fucosamine, α-D-glucosamine, and α-8-epi-legionaminic acid (13). Another study examined the sequences of an *A. baumannii* strain isolated from a fatal infection and showed that a plasmid that had evolved to harbor antibiotic resistance genes plays a role in the differentiation of cells specialized in the elimination of competing bacteria (18, 19). However, the genetic basis of the hypervirulent phenotype identified in a single hypervirulent strain may not represent a feature that is common in *A. baumannii*. In our study, the use of cgMLST to genotype the *A. baumannii* clinical isolates revealed a group of hypervirulent strains. Therefore, Lyb-2-4 appears to represent a common determinant of increased virulence, as was confirmed by finding it in virulent strains isolated in other institutions.

The structure of Lyb-2-4 in LAC-4 showed that the replacement of *orf4* and *orf6* by homologous genes in the CT512 Lyb-2-4 might not affect the hypervirulent phenotype, and suggests that Lyb-2-4 probably has different variants. Unfortunately, we don’t have the virulence information for the other strains we identified that also harbring Lyb-2-4: SAMN05601667, SAMN05601670, SAMN05362953, SAMN10411605, SAMN06650238 and SAMN05601671 (FIG. 4B). More studies are needed to investigate the correlation between the hypervirulent phenotype and the different variants of Lyb-2-4.

Because cgMLST queries thousands of alleles around the entire genome, it combines the comparability of MLST and the discriminatory power of pulsed-field gel electrophoresis (PFGE). It suggests that cgMLST is capable of uncovering new genomic determinants of phenotypic characteristics and therefore warrants widespread implemention.

Importantly, CRAB infections with isolates harboring Lyb-2-4 were associated with high mortality, highlighting the importance of rapidly detecting the presence of Lyb-2-4. The development of a PCR assay in this study should be very useful in advising the clinician to implement medical intervention earlier, and also making the worldwide surveillance of these hypervirulent CRAB strains easier. Implementation of strict control measures is needed to prevent these strains from further disseminating in hospital settings and also in the community.

## MATERIALS AND METHODS

### Bacterial isolates and phenotypic characterization

Consecutive non-replicate clinical isolates of CRAB isolated from patients in the ICU of Shenzhen People’s Hospital in 2017 were used in this study. Shenzhen people’s hospital is a medical center with 2500 beds in the Luohu district of Shenzhen, China. A VITEK-2 compact system (bioMérieux, Marcy-l’Étoile, France) was used to establish the strain identity and antimicrobial susceptibilities of the isolates. The results were interpreted in accordance with the guidelines published by the Clinical and Laboratory Standards Institute (CLSI; document M100-S26) (20). The species identity of all isolates was confirmed via matrix-assisted laser desorption/ionization mass spectrometry (bioMérieux, Marcy-l’Étoile, France).

### WGS and genomic characterization

Bacteria from frozen stocks were cultured in Luria-Bertani (LB) broth overnight at 37°C with shaking (220 rpm). Overnight cultures were diluted 1/100 and re-cultured until the optical density at 600 nm (OD_600_) was 0.6 – 0.8. Genomic DNA was extracted from each isolate using the Sodium Dodecyl Sulfate (SDS) method (21). The bacterial genomes were sequenced using Illumina HiSeq 2500 platform (Illumina, San Diego, CA, USA), and fastp was used to remove low-quality and low-complexity reads, and polyG/polyX tails (22). The genomes were assembled with de novo SPAdes Genome Assembler (version 3.12.0) (23).

Antimicrobial resistance genes and virulence factors in the isolates were identified by scanning the genome contigs against ResFinder and VFDB databases using ABRicate (version 0.8.7). We performed MLST and cgMLST genotyping and constructed minimum spanning trees (MST) with Ridom SeqSphere^+^ (version 5.1.0) (24). For *A. baumannii*, the cgMLST scheme (http://www.cgmlst.org/ncs/schema/3956907) consist of 2390 conserved genome-wide genes, and closely related genomes (≤9 alleles distance) were ‘lumped’ together as a complex type (CT).

### Transmission trees construction

We constructed transmission trees for the CT groups with more than 3 cases. Alignment and extraction of core genome from assembly contigs were performed with the Harvest suite (version 1.2) (25). Transmission trees were constructed based on the single nucleotide polymorphism (SNPs) in the core genome and integrated epidemiological information (i.e., patient exposure to infection) (Table S1) with BEAST2 (version 2.5.1) package Structured Coalescent Transmission Tree Inference (SCOTTI) (version 1.1.1) (26). SCOTTI that not only accounts for diversity and evolution within a host, but also for other sources of bias, namely non-sampled hosts and multiple infections of the same host. This new method builds on recent progress in efficiently modeling migration between populations using an approximation to the structured coalescent. SCOTTI only allowing hosts to transmit the disease during periods when they are infectious (between sampling and discharge date). We showed transmissions with a probability larger than 20%.

### Clinical data collection and genome comparison

Patient Clinical and epidemiological information were collected. To compare the genomes between CT512 and CT2085 group, we first extracted the conserved core genomes shared by CT512 isolates with Spine (version 0.3.2), and then used AGEnt (version 0.3.1) to compare the core genome of CT512 with the genomes of CT2085 to identify genes in the core CT512 genome absent from CT2085 (27). We annotated the results with Prokka (version 1.13.3) (28).

### Virulence test *in vivo*

We used a pneumonia model of *A. baumannii* in mice to test the virulence of isolates. C57BL/6 6-8 weeks old female mice under specific-pathogen-free grade were purchased from Hunan SJA laboratory animal CO., LTD (Hunan, China). The mice were intraperitoneally anaesthetized with pentobarbital sodium (75 mg/kg) and inoculated with 20 μL (2.5×10^8^, 5×10^7^, 1×10^7^, 0 CFU) of *A. baumannii* by non-invasive intratracheal instillation under direct vision. The survival of the mice was observed for 7 days post infection (8). All animal care and use protocols in this study were performed in accordance with the Regulations for the Administration of Affairs Concerning Experimental Animals approved by the State Council of People’s Republic of China. All animal experiments in this study were approved by the Animal Ethical and Experimental Committee of the Army Military Medical University (Chongqing, Permit No. 2011-04) in accordance with their rules and regulations.

### Rapid identification tool of Lyb-2-4

To develop a rapid tool to detect Lyb-2-4, all available *A. baumannii* NCBI completely integrated genome datasets (as of 2019-02-18) were downloaded. We compared the genomes with Lyb-2-4 to those without this locus and designed a polymerase chain reaction (PCR) assay to detect the presence of Lyb-2-4 in the genome. This PCR assay was then used to screen the isolates from 3 provinces of China, and the results were correlated with the mortality rate of each isolate.

### Data Availability

We deposited the genome sequences in GenBank under BioProject PRJNA533558.

## ACKNOWLEDGMENTS

This work was supported by the International Collaborative Research Fund (GJHZ20180413181716797) and Free Inquiry Fund (JCYJ20180305163929948) of Shenzhen Science and Technology Innovation Commission; We thank Yuting Huang for figures preparation.

## REFERENCES

1 Dijkshoorn L, Nemec A, Seifert H. An increasing threat in hospitals: multidrug-resistant *Acinetobacter baumannii*. Nature Rev Microbiol 2007; 5: 939–51.

2 Evans BA, Hamouda A, Amyes SG. The rise of carbapenem-resistant *Acinetobacter baumannii*. Curr Pharm Des 2013; 19: 223–38.

3 Giammanco A, Cala C, Fasciana T, et al. Global assessment of the activity of tigecycline against multidrug-resistant Gram-negative pathogens between 2004 and 2014 as part of the tigecycline evaluation and surveillance trial. mSphere 2017; 2: e00310–16.

4 Bertrand X, Dowzicky MJ. Antimicrobial susceptibility among gram-negative isolates collected from intensive care units in North America, Europe, the Asia-Pacific Rim, Latin America, the Middle East, and Africa between 2004 and 2009 as part of the Tigecycline Evaluation and Surveillance Trial. Clin Ther 2012; 34: 124–37.

5 WHO. Global priority list of antibiotic-resistant bacteria to guide researach, discovery and development of new antibiotics. World Health Organization, Geneva, 2017.

6 van Faassen H, Kuo Lee R, Harris G, Zhao X, Conlan JW, Chen W. Neutrophils play an important role in host resistance to respiratory infection with *Acinetobacter baumannii* in mice. Infect Immun 2007; 75: 5597–608.

7 Knapp S, Wieland CW, Florquin S, et al. Differential roles of CD14 and toll-like receptors 4 and 2 in murine Acinetobacter pneumonia. Am J Respir Crit Care Med 2006; 173: 122–9.

8 Zeng X, Gu H, Cheng Y, et al. A lethal pneumonia model of *Acinetobacter baumannii*: an investigation in immunocompetent mice. Clin Microbiol Infect 2019; 25: 516.

9 Harris G, Kuo Lee R, Lam CK, et al. A mouse model of Acinetobacter baumannii-associated pneumonia using a clinically isolated hypervirulent strain. Antimicrob Agents Chemother 2013; 57: 3601–13.

10 Ou HY, Kuang SN, He X, et al. Complete genome sequence of hypervirulent and outbreak-associated *Acinetobacter baumannii* LAC-4: epidemiology, resistance genetic determinants and potential virulence factors. Sci Rep 2015; 5: 8643.

11 Yu-Chung Chuang, Aristine Cheng, Hsin-Yun Sun, et al. Microbiological and Clinical Characteristics of *Acinetobacter baumannii* Bacteremia: Implications of Sequence type for Prognosis. J Infect 2019; 78:106–12.

12 Higgins PG, Prior K, Harmsen D, Seifert H. Development and evaluation of a core genome multilocus typing scheme for whole-genome sequence-based typing of *Acinetobacter baumannii*. PLoS ONE 2017; 12: e0179228.

13 Vinogradov E, Maclean L, Xu HH, Chen W. The structure of the polysaccharide isolated from *Acinetobacter baumannii* strain LAC-4. Carbohydr Res 2014; 390: 42–5.

14 Hu D, Liu B, Dijkshoorn L, Wang L, Reeves PR. Diversity in the major polysaccharide antigen of *Acinetobacter baumannii* assessed by DNA sequencing, and development of a molecular serotyping scheme. PLoS One 2013; 8: e70329.

15 Morrison MJ, Imperiali B. Biochemical Analysis and Structure Determination of Bacterial Acetyltransferases Responsible for the Biosynthesis of UDP-*N,N*’-Diacetylbacillosamine. J Biol Chem 2013; 288: 32248–60.

16 Russo TA, Luke NR, Beanan JM, et al. The K1 capsular polysaccharide of *Acinetobacter baumannii* strain 307-0294 is a major virulence factor. Infect Immun 2010; 78: 3993–4000.

17 Hacker J, Kaper JB. Pathogenicity islands and the evolution of microbes. Annu Rev Microbiol 2000; 54: 641–79.

18 Ahmed-Bentley J, Chandran AU, Joffe AM, French D, Peirano G, Pitout JD. Gram-negative bacteria that produce carbapenemases causing death attributed to recent foreign hospitalization. Antimicrob Agents Chemother 2013; 57: 3085–91.

19 Weber BS, Ly PM, Irwin JN, Pukatzki S, Feldman MF. A multidrug resistance plasmid contains the molecular switch for type VI secretion in *Acinetobacter baumannii*. Proc Natl Acad Sci U S A 2015; 112: 9442–7.

20 CLSI. Performance standards for antimicrobial susceptibility testing; twenty-sixth informational supplement. CLSI document M100-S26. Clinical and Laboratory Standards Institute: Wayne, PA, 2016.

21 Chen WP, Kuo TT. A simple and rapid method for the preparation of gram-negative bacterial genomic DNA. Nucleic Acids Res 1993; 21: 2260.

22 Chen S, Yanqing Z, Yaru C, Jia G. fastp: an ultra-fast all-in-one FASTQ preprocessor. Bioinformatics 2018; 34: 884–90.

23 Antipov D, Korobeynikov A, McLean JS, Pevzner PA. hybridSPAdes: an algorithm for hybrid assembly of short and long reads. Bioinformatics 2016; 32: 1009–15.

24 Jünemann S, Sedlazeck FJ, Prior K, et al. Updating benchtop sequencing performance comparison. Nat Biotechnol 2013; 31: 294–6.

25 Treangen TJ, Ondov BD, Koren S, Phillippy AM. The Harvest suite for rapid core-genome alignment and visualization of thousands of intraspecific microbial genomes. Genome Biol 2014; 15: 524.

26 De Maio N, Wu CH, Wilson DJ. SCOTTI: Efficient Reconstruction of Transmission within Outbreaks with the StructuredCoalescent. PLoS Comput Biol 2016; 12: e1005130.

27 Ozer EA, Allen JP, Hauser AR. Characterization of the core and accessory genomes of *Pseudomonas aeruginosa* using bioinformatic tools Spine and AGEnt. BMC Genomics 2014; 15: 737.

28 Seemann T. Prokka: rapid prokaryotic genome annotation. Bioinformatics 2014; 30: 2068–9.

